# Denoising-based Image Compression for Connectomics

**DOI:** 10.1101/2021.05.29.445828

**Authors:** David Minnen, Michał Januszewski, Tim Blakely, Alexander Shapson-Coe, Richard L. Schalek, Johannes Ballé, Jeff W. Lichtman, Viren Jain

## Abstract

Connectomic reconstruction of neural circuits relies on nanometer resolution microscopy which produces on the order of a petabyte of imagery for each cubic millimeter of brain tissue. The cost of storing such data is a significant barrier to broadening the use of connectomic approaches and scaling to even larger volumes. We present an image compression approach that uses machine learning-based denoising and standard image codecs to compress raw electron microscopy imagery of neuropil up to 17-fold with negligible loss of 3d reconstruction and synaptic detection accuracy.

## Main Text

Progress in the reconstruction and analysis of neural circuits has recently been accelerated by advances in 3d volume electron microscopy (EM) and automated methods for neuron segmentation and annotation^1,2^. In particular, contemporary serial section EM approaches are capable of imaging hundreds of megapixels per second^3^,^4^, which has led to the successful acquisition of petabyte-scale volumes of human^5^ and mouse^6^ cortex. However, scaling further poses substantial challenges to computational infrastructure; nanometer resolution imaging of a whole mouse brain^7^ could, for example, produce one million terabytes (an exabyte) of data. While storage systems with such capacities are available, they are costly to build and operate. More generally, even petabyte-scale requirements limit the number and diversity of circuit reconstruction efforts. Thus in order to mitigate the costly storage demands of connectomics, we developed a practical approach to compression of electron microscopy data that yields at least a 17-fold reduction in storage requirements.

The key insight is that computational removal of imaging noise allows EM images to be compressed far more effectively than in their raw form^8,9^. Since an effective noise model for complex microscopy images can be difficult to formulate *a priori*, we instead learn a denoising model directly from EM imagery itself. We acquired pairs of EM scans of human cortical tissue^5^ under different imaging speeds of a Zeiss 61-beam scanning electron microscope^3^, which resulted in images of the same tissue but containing different amounts of noise (i.e., fast with more noise vs slow with less noise). After precise alignment of fast and slow scans (see Methods), the image pairs were used to train a neural network^10^ with standard supervised techniques. The trained network was then used to infer “slow” 3200 nanosecond low-noise scans (Figure 1) from the “fast” 200 nanosecond acquisition scans (acquiring all data in the low-noise regime would be too slow for large tissue volumes which require months of imaging^5,6)^.

**Figure 1.**
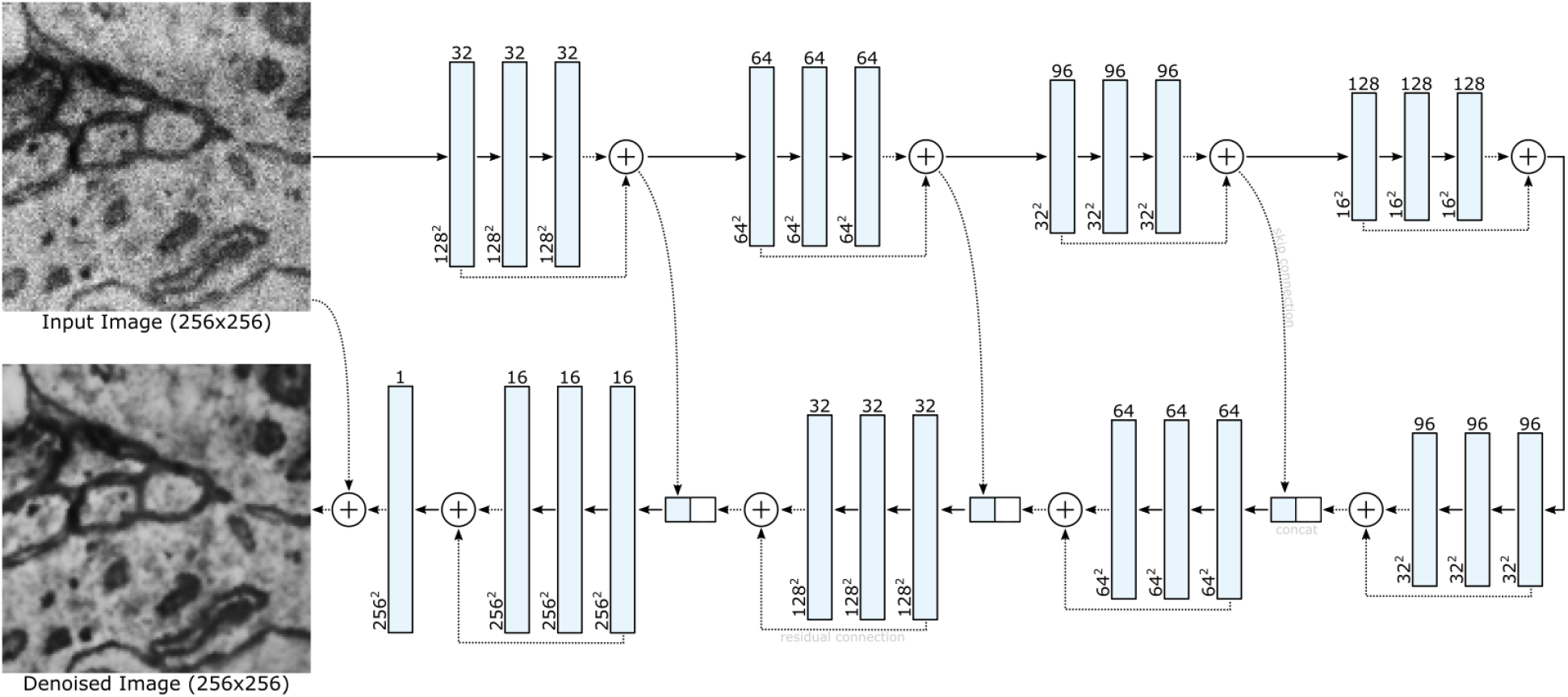
A deep neural network learns to denoise tissue scans after being optimized on aligned pairs of high-noise (fast dwell time) and low-noise (slow dwell time) scans. For efficient processing, the network uses a U-Net architecture^10^ with residual connections for both the individual blocks and for the final output.

Denoised EM imagery was subsequently compressed using lossy codecs (JPEG XL or AVIF). These codecs provide tunable parameters that trade off compression rate for image fidelity. Beyond visual inspection (Figure 2) we wanted to establish that automated 3d reconstruction of the data would not be degraded by denoising or compression. Therefore we used flood-filling networks (FFNs)^11^ to segment neurons based on original, compressed, and denoised+compressed data and compared the reconstruction results to human ground truth (see Methods).

**Figure 2.**
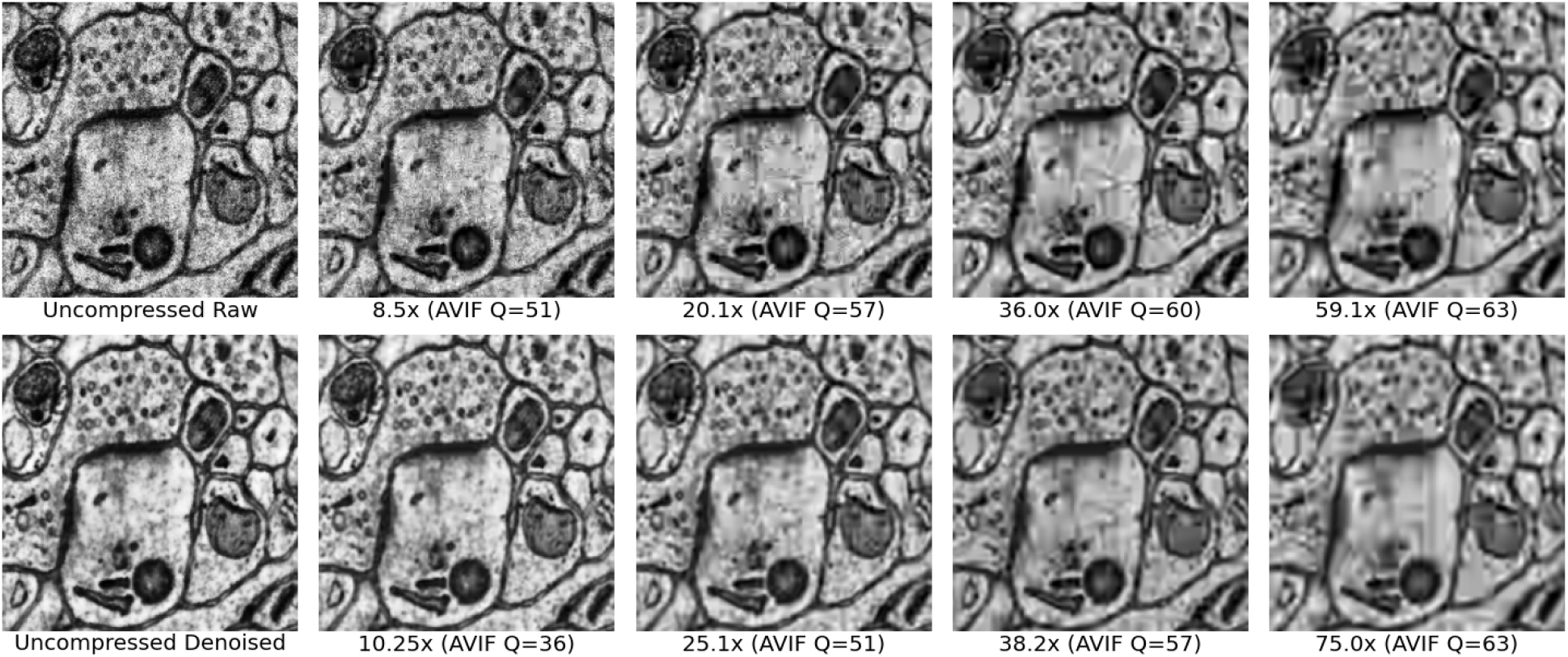
By first denoising the raw data, both compression rates and visual quality increase (bottom row) compared to directly compressing the noisy scans (top row). Specifically, fewer block boundary artifacts (due to the block-based partition used by AVIF) and less ringing (due to the use of discrete cosine or wavelet transforms) appear in the compressed version of the denoised scans. Online side-by-side views at comparable compression rates are available for **AVIF** and **JXL**.

We found that denoising alone did not impact reconstruction accuracy and that compression of denoised data enabled 17-fold storage reduction with negligible loss in reconstruction quality according to an “edge accuracy” metric that quantified the number of splits and mergers in an automated result^11^ (Figure 3). This compression was more than 3x greater compared to what was achievable with data that was compressed *without* first being denoised.

**Figure 3.**
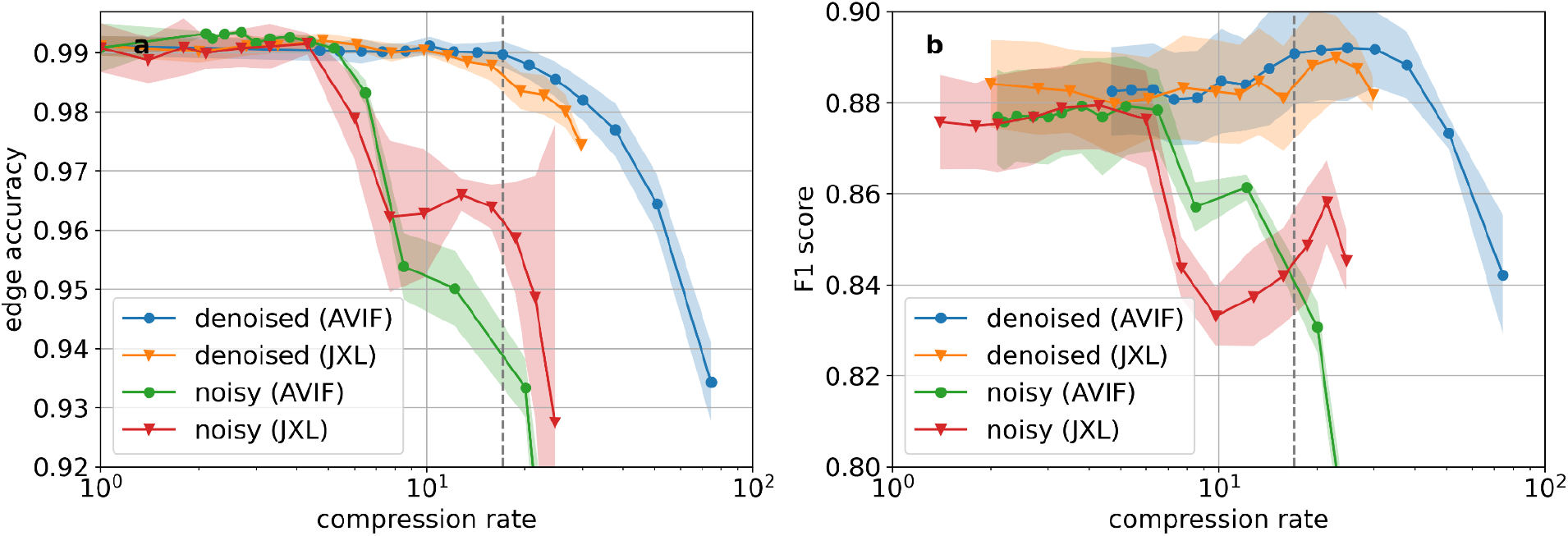
a) Segmentation quality of noisy and denoised images as a function of compression rate. Points and lines show the means, and the shaded area covers ±1 standard deviation around the mean. Statistics were computed over segmentations of a 9.56 · 10^4^ μm^3^ region, generated with 5 different network weight snapshots (“checkpoints”) of the FFN operating on 8×8×30 nm^3^ imagery. The dashed line corresponds to a 17.1x compression rate. b) Synapse detection performance. Statistics are computed over 5 different model checkpoints. Compression rates are computed with 8×8×30 nm^3^ voxel volume as the baseline, which was the highest resolution used for automated processing. Taking downsampling into account, 17.1x therefore corresponds to a 68.4x size reduction relative to the size of images as originally acquired with 4×4 nm^2^ pixels.

In addition to 3d reconstruction, connectomic analysis requires identification of synapses. We therefore performed synaptic detection on a 19.2×19.2×9.93 μm volume of human cortex data using a U-Net trained on human ground truth and compared detection results on original, compressed, and denoised+compressed data. As with FFN segmentation, we found that denoising did not affect detection results and that compression which achieves 17-fold storage reduction had a negligible effect on synapse detection (Figure 3).

We believe the proposed approach is a highly practical method of achieving compression in real-world connectomics pipelines, for several reasons. First, training the denoising model requires an insignificant amount of additional imaging relative to an overall large-scale acquisition effort. For instance, in our experiments we used tens of gigabytes of images acquired at slow and fast acquisition speeds, which was equivalent to less than 0.005% of the complete dataset. Second, both the denoising and compression codecs operate on 2d data which means the models can be applied to unstitched and unaligned raw data directly from the microscope. Third, denoising involves inference with a relatively standard U-Net neural network architecture for which inference can be computed at tens of megapixels per second on a single GPU^12^; compression involves codecs such as JPEG XL which also can be computed at tens of megapixels per second on a single CPU^13^.

Finally, we note that our approach does not rely on a learned compression model^14^. This simplifies implementation and enables a flexible approach to choosing compression parameters. That said, learned compression remains an interesting avenue for potential further reductions in storage requirements. The application of volumetric learned compression models^15^ to post-aligned EM data is also an interesting research direction. However, the approach described here already reduces storage requirements by an order of magnitude and thus dramatically increases feasibility of existing and proposed^7^ projects.

## Supplementary Methods

### Electron Microscopy Data

We used a small subset of the 1 mm^3^ human cortex H01 sample^5^ for all experiments. The surgical tissue was extracted from the left medial temporal lobe of a 45-year old woman with a history of seizures, and fixed immediately after excision. The excised sample was free of gross pathology. It was subsequently stained using the ROTO protocol, embedded in resin, and sectioned with an ATUM device. The part of the sample we used was cut with a section thickness of 30 nm. The sections were affixed to silicon wafers and imaged with a 61-beam Zeiss MultiSEM microscope at a pixel resolution of 4×4 nm^2^, and with a beam dwell time of 200 ns, current of 576 pA, and landing energy of 1.5 kV.

The image tiles for this project were stitched and aligned by Adi Peleg using a custom software implementation of elastic alignment^16^. We normalized the contrast of the aligned volume with contrast limited adaptive histogram equalization (CLAHE) applied grid-wise within 1300×1300-pixel patches with an overlap of 300×300 pixels.

### Scan Registration

Training data is extracted from two sets of scans of the same tissue, one at 200 ns and one at 3200 ns. The corresponding images between the two scans have considerable overlap but are not precisely aligned as captured. To achieve pixel-accurate alignment, each 3200 ns image is processed by a four-stage pipeline. First, all images are normalized using CLAHE. Second, each 3200 ns image is roughly aligned with its 200 ns counterpart via translation by estimating the 2d offset between the two images using cross-correlation. The grayscale pixel values of the 3200 ns image are then adjusted to better match the 200 ns image using a linear transform. In this step, we solve a linear regression problem to find the best scalar and additive term to minimize the squared error between each grayscale value in the roughly aligned 3200 ns and the corresponding pixel in the 200 ns image. Finally, the image undergoes nonrigid warping according to a flow field estimated by multiscale TV-L1 optical flow^17^.

### Denoising

The denoising model works by learning to map noisy scans captured using 200 ns dwell times to low-noise scans captured using 3200 ns dwell times. The network architecture follows the U-Net approach^10^, where input images are analyzed at successively lower resolutions down one path and then mapped to output images via upscaling on a second path (Figure 1). Skip connections between the two paths allow the network to more easily utilize high-resolution information, while residual connections are used both within each block in the U-Net and for the final output.

Our model uses U-Net blocks with three convolutional layers. The first uses a stride of two with 5×5 kernels, while the second and third layers use a stride of one with 3×3 kernels. In the upscaling path, the stride two convolutions are replaced with transposed convolution to increase, rather than decrease the spatial resolution, and all layers use “same” padding. Gaussian Error Linear Units (GELUs)^18^ are used between each convolutional layer.

Four residual blocks are used in the downscaling path to reduce the resolution by a total of 16x. The blocks increase the channel depth from a single grayscale input image to 32, 64, 96, and finally 128 channels. The upscaling path mirrors this structure and uses depths of 96, 64, 32, and 16 for its blocks. A single final convolution using a 3×3 kernel with an output depth of one generates a grayscale residual. The denoised image is then produced by adding this residual to the input image.

Training makes use of 256×256 patch pairs extracted from the aligned scans. The network is fully convolutional so the input resolution is not explicitly constrained by the architecture, but limited memory on real hardware requires us to tile the input for very large images. When denoising on a CPU, we used 4096×4096 tiles, but smaller tiles may be required for GPUs with less onboard memory. To minimize boundary artifacts, we used an additional border of 128 pixels on all sides that is dropped after denoising. In the case where 128 extra pixels are not available, e.g., at the edge of the scan volume, reflection-based padding is used.

To drive the optimization process, a differentiable loss function is needed. We do not expect perfect prediction of the low-noise scans, and so, ideally, the loss function would optimally score image discrepancies in proportion to their effect on downstream analysis both by scientists and by automated systems. In theory, we could use the segmentation algorithm as a loss function, but it is currently a multi-stage process that is not fully differentiable. Regardless, even if we could train in an end-to-end fashion with the segmentation algorithm, the implied image quality metric may not preserve details important to scientists.

In the absence of a task-specific loss, we used a mixture of three common image quality metrics: L1 error, structural similarity (SSIM)^19^, and the Learned Perceptual Image Patch Similarity (LPIPS) metric^20^. We empirically settled on weights of 0.2, 0.5, and 0.3 for the three metrics, respectively.

The model was trained for 600,000 steps using a batch size of eight and the Adam^21^ optimizer. The learning rate was initially set to 10^-4^ and linearly reduced to 10^-5^ between steps 350k and 550k, and then down to 10^-6^ by step 600k. We experimented with other nonlinear activation functions (relu, leaky relu, swish, tanh, sigmoid, etc.), different normalization layers (instance, layer, and channel-normalization), and with different weights for the image metrics. Visually, changing the activation function did not have a large effect, normalization layers degraded visual quality, and the loss weights only had a significant effect when extreme values were used, e.g., a weight of 1.0 for LPIPS and 0.0 for L1 and SSIM.

### Codecs

We experimented with two state-of-the-art codecs: JPEG XL (JXL) and the AV1 Image File Format (AVIF). Both are suitable for our task and outperform older image codecs such as JPEG, JPEG2000, and WebP. Either codec could be used in practice, with JPEG XL showing faster encode/decode times, while AV1 supports higher compression rates and slightly better rate-distortion performance. We believe the compression quality difference is because JPEG XL was optimized for near lossless subjective quality, which may not transfer well to the domain of EM images of brain tissue. We generated images at different compression rates by varying the JXL distance and AVIF *q* parameter. We set the JXL effort option to 9 and the AVIF speed parameter to 1. We observed that denoising led to higher relative gains in compression rates when the AVIF codec was used (see Sup. Fig. 1).

**Sup. Fig. 1.**
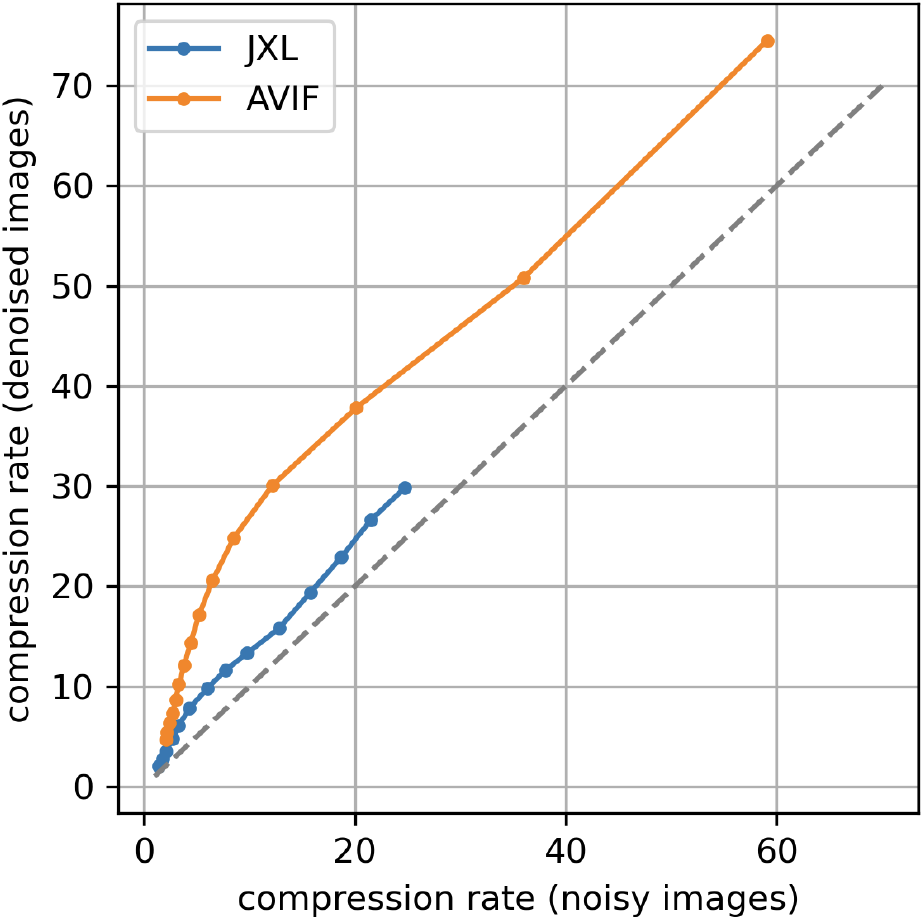
Compression rate of noisy and denoised images at constant JXL distance and AVIF *q* parameter.

### Flood-filling Network Segmentation and 3d Reconstruction Evaluation

We performed automated reconstruction experiments over a subset of the H01 volume (“ROI215”) — a 98.3 × 98.3 × 9.9 μm^3^ region within layer 2 of the cortex. For neuron segmentation we used a simplified version of the multiscale flood-filling network (FFN) pipeline^5^,^11^. Specifically, we trained 6 separate FFN models operating on original or denoised data at 32×32×30 nm^3^, 16×16×30 nm^3^ and 8×8×30 nm^3^ voxel size. We denoised the images at their original acquisition resolution of 4×4×30 nm^3^ per voxel and downsampled them with area-averaging to 8×8×30 nm^3^ voxel size. Compression, when used, was applied at 8×8×30 nm^3^ resolution, before further downsampling. The compression rates we report use the 8×8×30 nm^3^ image sizes as baseline. We used the FFN models to first build a “base segmentation” minimizing the number of false mergers, and then applied an agglomeration procedure to reduce split errors.

We computed the oversegmentation consensus^11^ between four input segmentations — the outputs of the 32×32×30 nm^3^ and 16×16×30 nm^3^ FFN models each executed with forward and reverse seed ordering, and filtered to retain segments comprising 10,000 voxels or larger at their respective resolution. The base segmentation was then formed by filling the remaining unsegmented voxels with the 8×8×30 nm^3^ FFN model. Unlike the full H01 dataset, ROI215 does not contain any sufficiently large image irregularities (alignment problems, imaging artifacts) to require the use of FFN movement restriction or tissue masking.

We used FFN resegmentation to agglomerate objects from the base segmentation as described previously^11^. We computed agglomeration scores with all three FFN models (for 32×32×30 nm^3^, 16×16×30 nm^3^ and 8×8×30 nm^3^ voxel sizes), and considered a segment pair as an edge in the agglomeration graph if the scores fulfilled the agglomeration criteria11 in the results of any of the three FFN models. For the agglomeration we only considered pairs of segments: 1) present prior to the 8×8×30 nm^3^ fill-in when using the 16×16×30 nm^3^ and 32×32×30 nm^3^ FFN models; and 2) not already evaluated in step 1) when using the 8×8×30 nm^3^ FFN model. Every agglomeration graph was post-processed to enforce separation between the 29 somas contained within the volume^5^.

We compared the 50 Gvx FFN segmentations to a proofread sparse skeleton reconstruction of ROI215 with 0.97M skeleton edges covering 14,568 neurite fragments with a total path length of 282 mm (Sup. Fig. 2). For every segment, we classified the corresponding skeleton edges as correct, split, merged, or omitted, and from these classifications derived the edge accuracy and expected run length (ERL) metrics^11^ to quantify how well the automated segmentation reflected the ground truth skeletons. We randomly split the ground truth skeletons 19:1 into a testing and a tuning set. The maximum ERL achievable with the testing skeleton set was 35 μm.

**Sup. Fig. 2.**
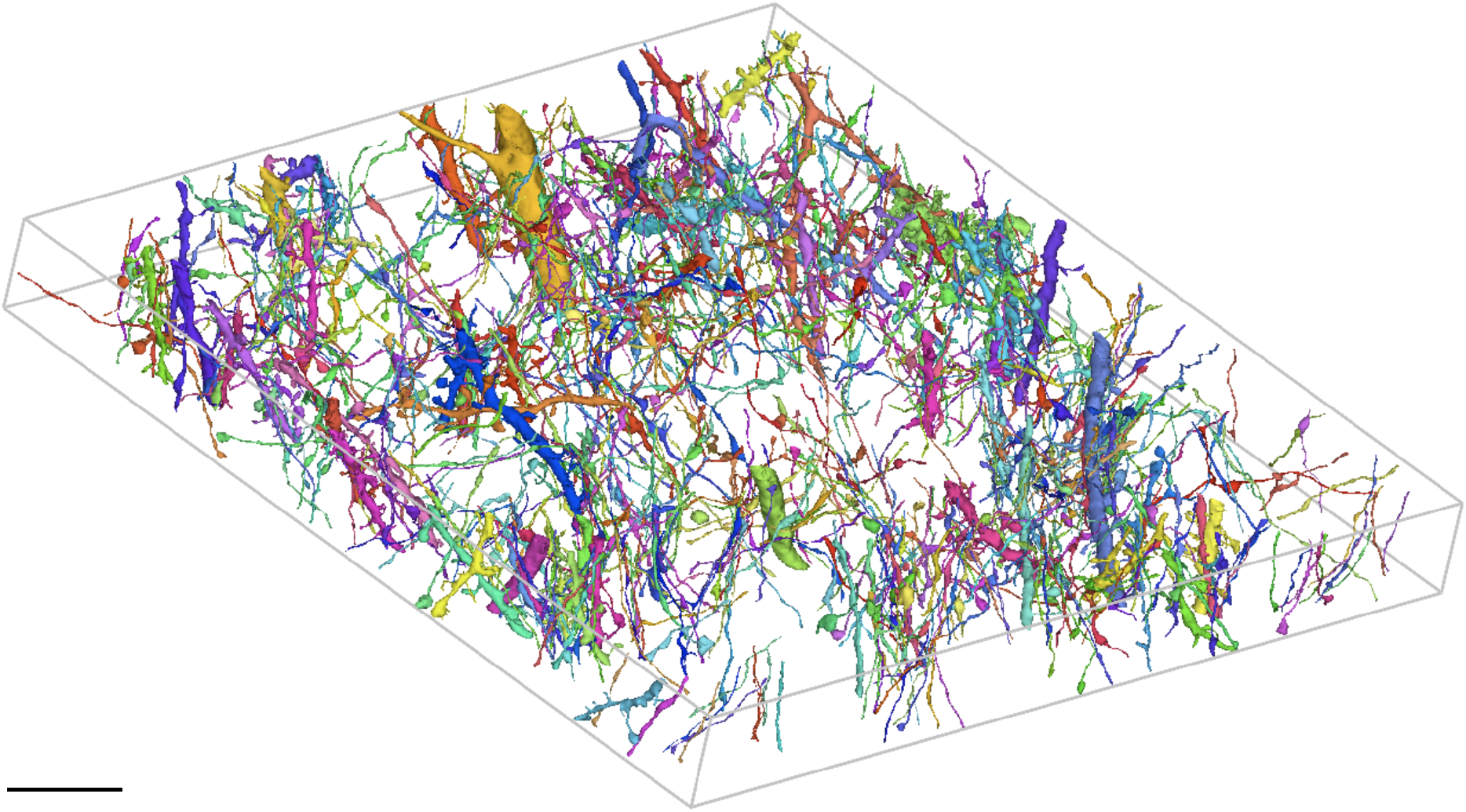
Random 5% sample of neurites in the ground truth set. Scale bar: 10 μm.

We found that the lowest resolution (32×32×30 nm^3^) FFN model trained on noisy images could be applied to denoised images without loss of accuracy, but the same was not true for higher resolution models (Sup. Fig. 3). In segmentation experiments we therefore always used segmentation models exclusively with the type of images (noisy, denoised) they were trained on.

**Sup. Fig. 3.**
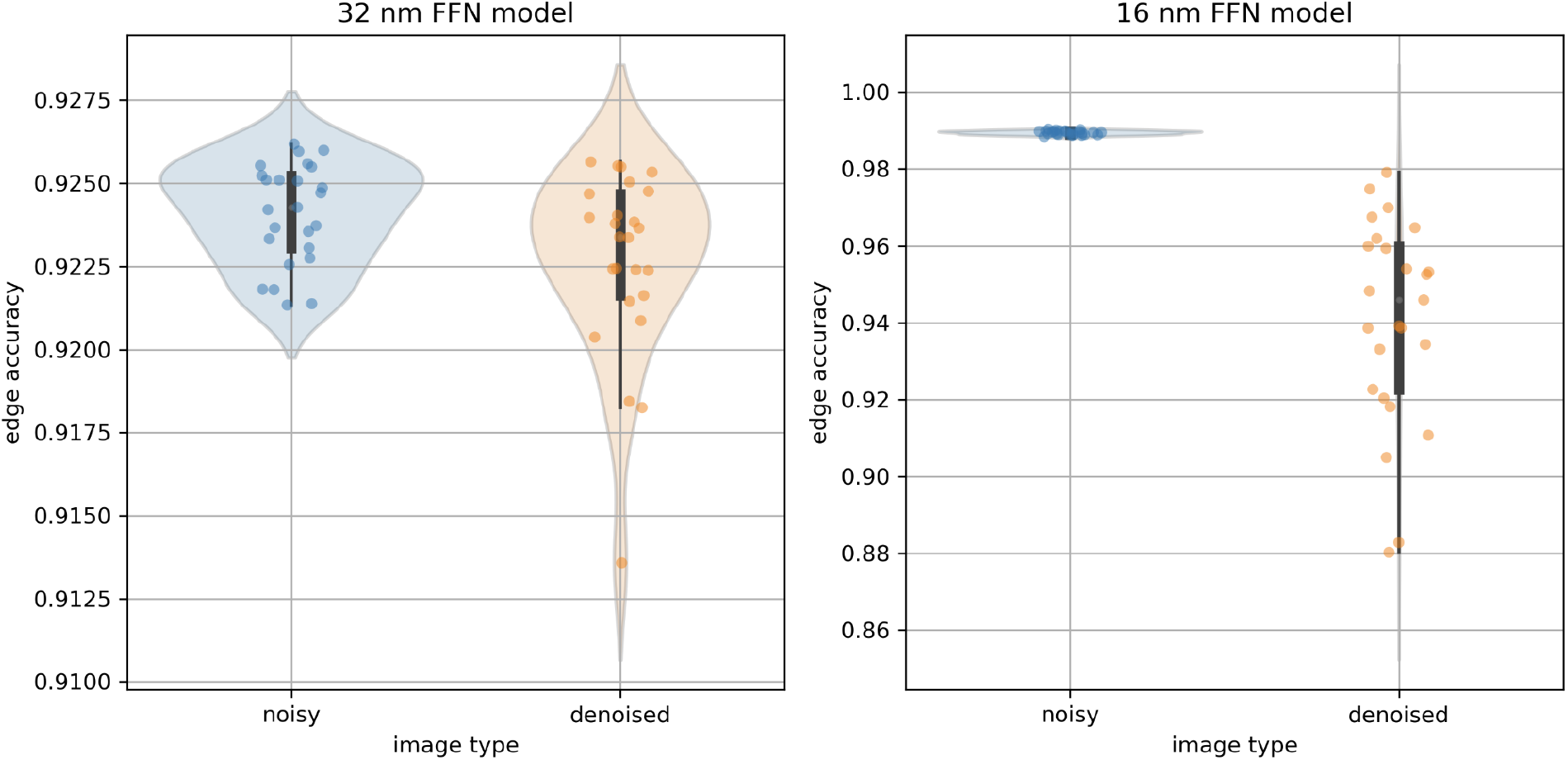
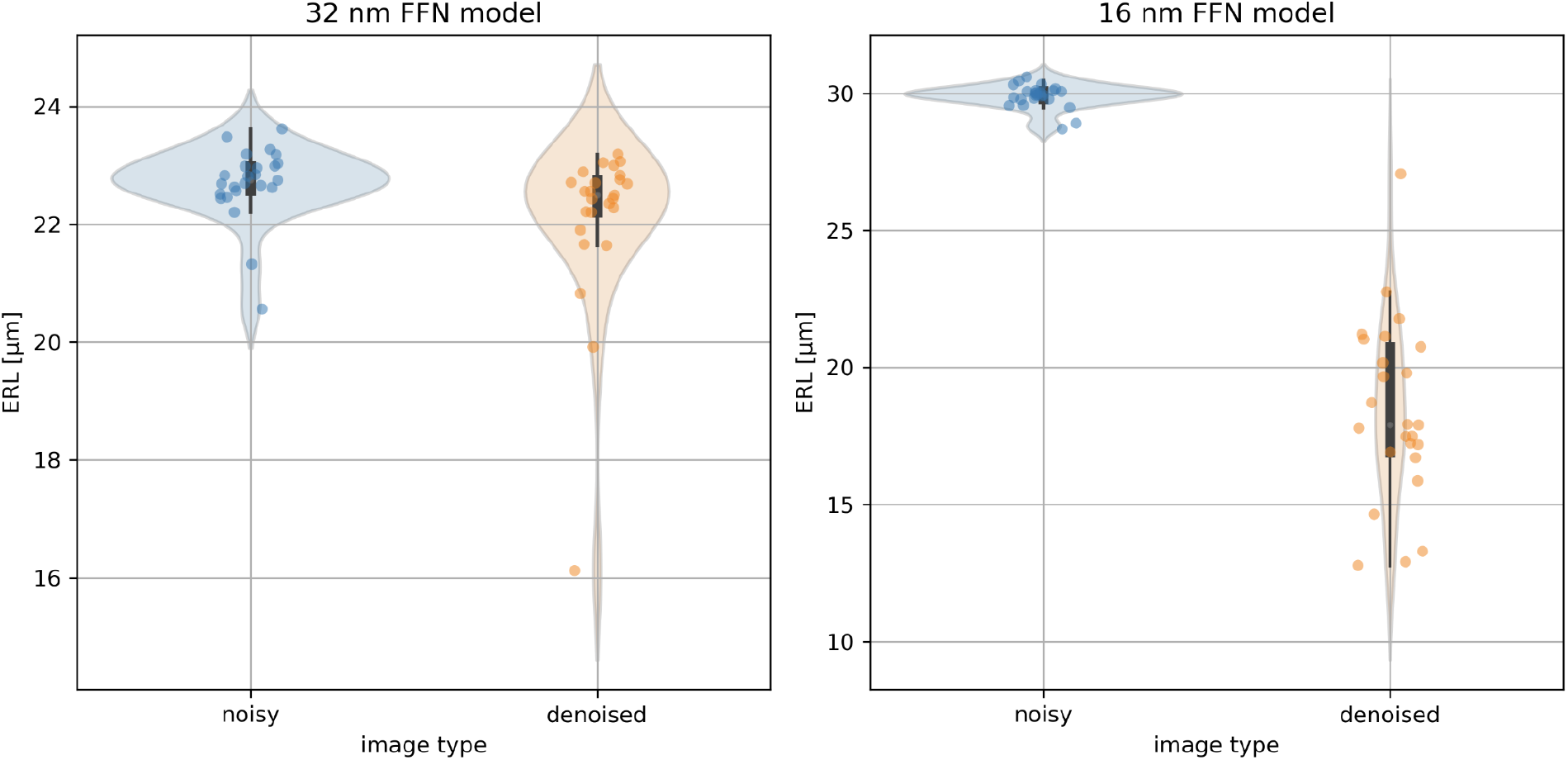
Impact of image denoising on segmentation models trained on noisy data only. Dots represent weight snapshots (checkpoints) of the respective FFN model. All results in the main text were obtained by applying models to the type of data (noisy, denoised) they were trained on.

**Sup. Fig. 4.**
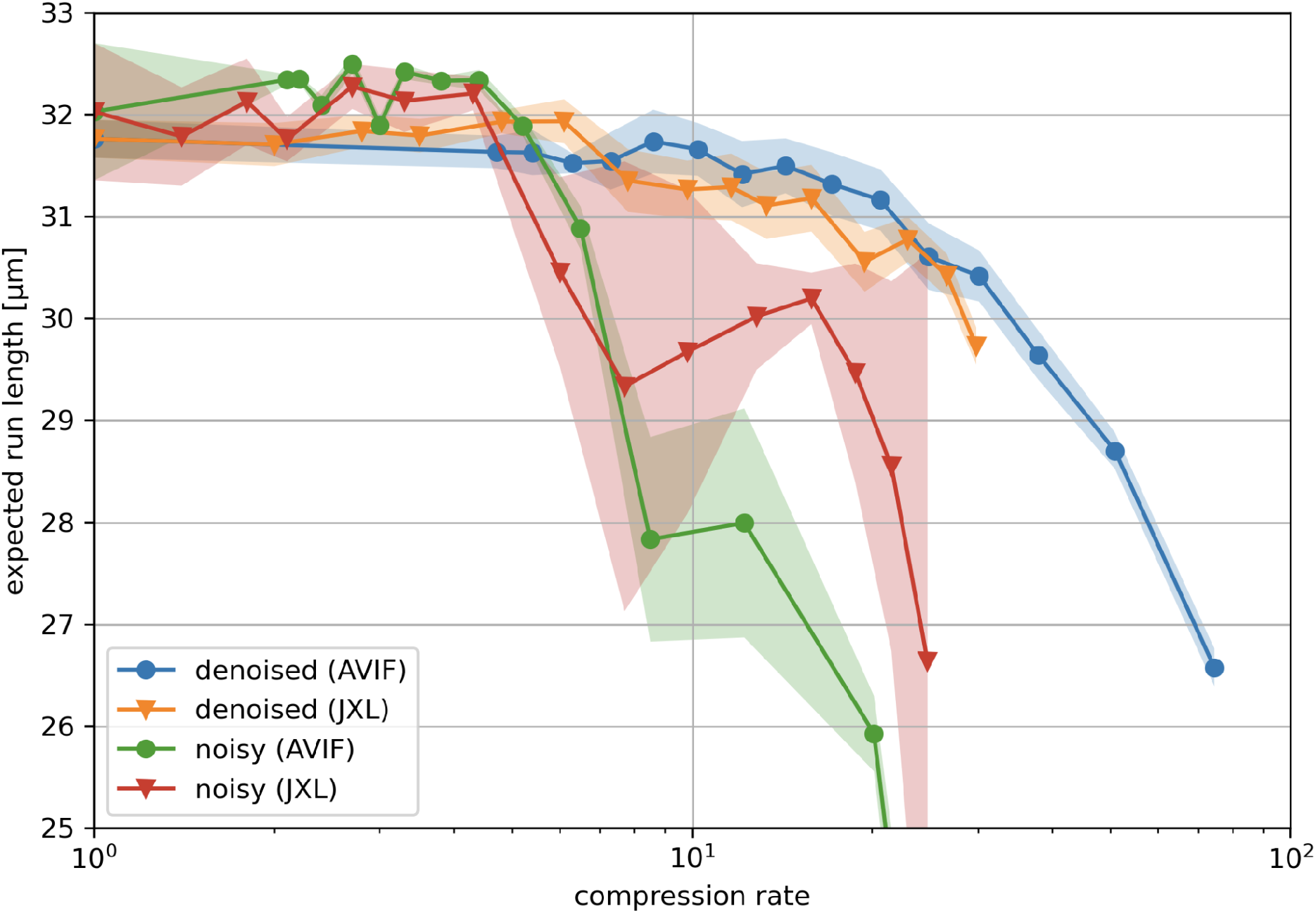
Segmentation quality quantified with ERL for compressed noisy and denoised images. Note that relative quality trends closely track those seen in Fig. 3.

### Synaptic Site Detection and Synapse Assembly

We performed automated synapse detection over the same subset of H01 (“ROI215”) that was used for segmentation experiments. Voxel-wise inference was performed to classify pre-synaptic, post-synaptic, and background using a previously described U-Net style model ^5,10^. Ground truth for the model was generated by densely labeling 6 ROIs across six cortical layers, each a subvolume of 600×600×3600 voxels (4.8 × 4.8 × 118.8 μm). Pre- and postsynaptic labels were proofread for correctness, resulting in a total of 3,824 synaptic examples. Synaptic site models were trained on noisy and denoised versions of the six ROIs, and inferred on noisy and denoised versions of the ROI215 dataset, respectively.

We assembled nearby pre- and postsynaptic sites into unique synapses as previously described^5^. Connected components were computed over the volume to assign each pre-/post-synaptic site a unique ID. Synaptic sites were then split according to the segmentation, where any potential site that spanned more than one segment and had more than 20% of its voxels within a given overlapping segment was split into two or more unique IDs and re-labeled. Synaptic sites were then filtered, enforcing a minimum volume of 100 voxels and removing any site in which more than 20% of the voxels were overlapping blood vessels or myelination according to the tissue masking. Next, a k-nearest-neighbor search within a radius of 800 nm was performed to identify potential pairs of pre-/post-sites. The search included both pre- and postsynaptic sites to allow for potentially polyadic synaptic connections. Furthermore, pairs were discarded if both sites contained the same underlying segment ID, or at least one synaptic site had the majority of its voxels overlapping un-segmented (background) data in the segmentation.

A final post-processing step was done to ensure that redundantly identified synapses were merged into a single connection (this can occur if there are multiple distinct postsynaptic densities for a single synapse). Synapses overlapping the same pre/post segment ID were grouped together, and if any two synapses were within 1 μm they were considered redundant identifications and the volumetrically smaller synapse was dropped.

We evaluated the impact of denoising and compression by comparing synapse detections to a proofread subset of 691 synapses within ROI215 that were deemed to be chemical in nature, either excitatory or inhibitory. Each ground truth synapse was considered recovered if there was a predicted synapse whose center of mass was within 250 nm. We verified that >99% of unique synapses in the ground truth set were >= 250 nm apart. The performance of the synapse models closely followed the performance of the segmentation models, with no significant differences from baseline accuracy up to or beyond the 17.1x compression rate for both codecs.

To disentangle the impact of the segmentation on the overall synapse detection pipeline we also evaluated all synapse predictions postprocessed with the same proofread segmentation volume (see Sup. Fig. 5). This revealed that while the F_1_ score remained close to the baseline level up to 30x compression on denoised images, recall started to degrade already around the 17.1x compression rate.

**Sup. Fig. 5.**
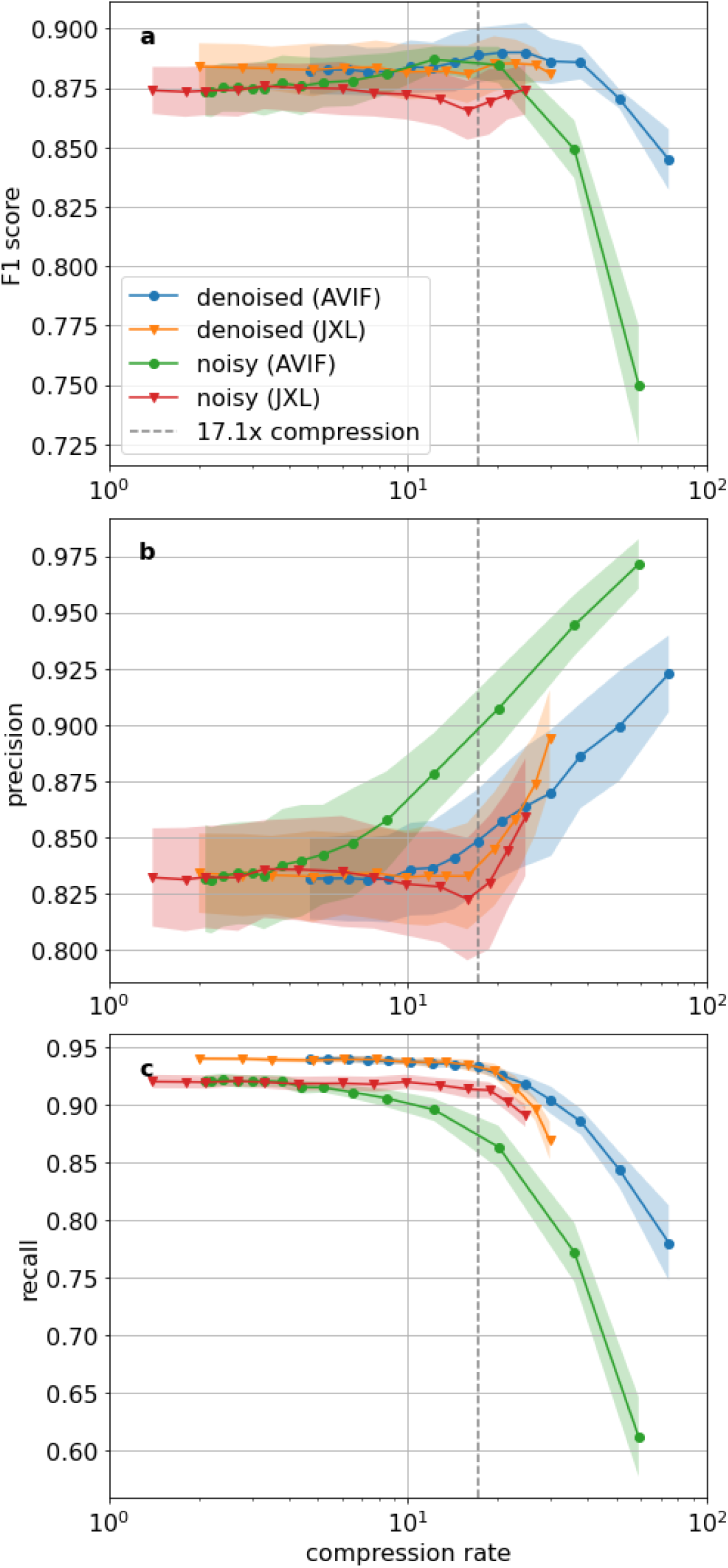
Synapse detection performance independent of segmentation (all predictions were postprocessed with the same proofread segmentation) quantified by a) F_1_ score, b) precision, c) recall.

